# MAPS: model-based analysis of long-range chromatin interactions from PLAC-seq and HiChIP experiments

**DOI:** 10.1101/411835

**Authors:** Ivan Juric, Miao Yu, Armen Abnousi, Ramya Raviram, Rongxin Fang, Yuan Zhao, Yanxiao Zhang, Yuchen Yang, Yun Li, Bing Ren, Ming Hu

## Abstract

Hi-C and chromatin immunoprecipitation (ChIP) have been combined to identify long-range chromatin interactions genome-wide at reduced cost and enhanced resolution, but extracting the information from the resulting datasets has been challenging. Here we describe a computational method, MAPS, Model-based Analysis of PLAC-seq and HiChIP, to process the data from such experiments and identify long-range chromatin interactions. MAPS adopts a zero-truncated Poisson regression framework to explicitly remove systematic biases in the PLAC-seq and HiChIP datasets, and then uses the normalized chromatin contact frequencies to identify significant chromatin interactions anchored at genomic regions bound by the protein of interest. MAPS shows superior performance over existing software tools in analysis of chromatin interactions centered on cohesin, CTCF and H3K4me3 associated regions in multiple cell types. MAPS is freely available at https://github.com/ijuric/MAPS.

## Main text

While millions of candidate enhancers have been predicted in the human genome, annotation of their target genes remains challenging, because these elements are known to regulate genes far away and often not their immediate neighbors^1^. Mapping of long-range chromatin interactions between enhancers and target gene promoters has been increasingly used to predict the target genes of enhancers and dissect gene regulatory networks^2^. Chromosome conformation capture (3C)^3^ based methods, such as *in situ* Hi-C^4^, have been used to detect long-range chromatin interactions in mammalian cells. However, billions of reads are typically needed to achieve kilobase resolution, limiting their applications. PLAC-seq^5^ and HiChIP^6^ technologies combine *in situ* Hi-C and chromatin immunoprecipitation (ChIP) to efficiently capture chromatin interactions anchored at genomic regions bound by specific proteins or histone modifications, achieving kilobase (Kb) resolution with much reduced sequencing cost^7^. However, compared to *in situ* Hi-C, the ChIP procedure introduces additional layers of experimental biases, posing significant challenges for data analysis.

Several software tools, including Fit-Hi-C^8^, HiCCUPS^4^, Mango^9^ and hichipper^10^ have been used to process PLAC-seq and HiChIP data. However, these methods are not optimal for such datasets. Specifically, both Fit-Hi-C and HiCCUPS are developed to analyze Hi-C data, and do not take into account the experimental biases introduced by the ChIP procedure. Mango is developed for ChIA-PET, a method that is similar to PLAC-seq and HiChIP, but carries different set of experimental biases due to the reverse order of the ChIP and proximity ligation procedures. Further, it only assesses the chromatin interactions between genomic regions bound by the protein of interest. In practice, chromatin interactions may also occur between a genomic region bound by the protein and other unoccupied regions, and Mango would miss such interactions due to the use of the filter. Hichipper^10^ is tailored for HiChIP data but still relies on Mango to identify long-range chromatin interactions, thus suffers the same weakness (more discussions in ***Supplementary Note 1***).

MAPS addresses the above issues by taking into account of the unique experimental setup and biases of PLAC-seq and HiChIP data in each step of data pre-processing, normalization and chromatin interaction determination (***Figure 1a***). First, MAPS splits the intra-chromosomal reads into two groups: short-range reads (<=1Kb) corresponding to the primary sites of protein occupancy and long-range reads (>1Kb) informative of the captured long-range chromatin interactions (***Figure S1***). Each valid long-range paired-end read is further assigned into “NOT”, “XOR” or “AND” set of bin pairs in the contact matrix depending on whether the pair of bins is bound by the protein of interest. As illustrated in ***Figure 1a***, the “NOT” set refers to bin pairs not overlapping any ChIP-seq peaks; the “XOR” set refers to bin pairs with only one end overlapping ChIP-seq peaks; whereas the “AND” set refers to those with both bins overlapping the ChIP-seq peaks. MAPS models interactions in the “AND” and “XOR” sets only, since bin pairs in the “NOT” set are not targets of the experiment and often contain very few reads for a reliable interaction call. Second, MAPS considers a variety of systematic biases shared by all 3C-based methods including differential effective fragment length, GC content, and sequence uniqueness (i.e., mappability)^11^, and additional experimental noises introduced by the ChIP procedure in PLAC-seq and HiChIP experiment (***Figure S2***). To mitigate these systematic biases, MAPS incorporates a modified version of HiCNorm^12^ algorithm, which successfully removes most of the biases, as illustrated by greatly reduced Pearson correlation coefficients between contact frequency and each bias factor (***Figure S2***). MAPS then calculates the expected contact frequency, P-value and false discovery rate (FDR) of each bin pair in “XOR” and “AND” sets. Noticeably, MAPS treats “XOR” or “AND” sets as two independent groups for data normalization, since bin pairs in the “AND” set have much higher chromatin contact frequency than bin pairs in the “XOR” set due to ChIP enrichment on both ends (***Figure S3***).

**Figure 1.**
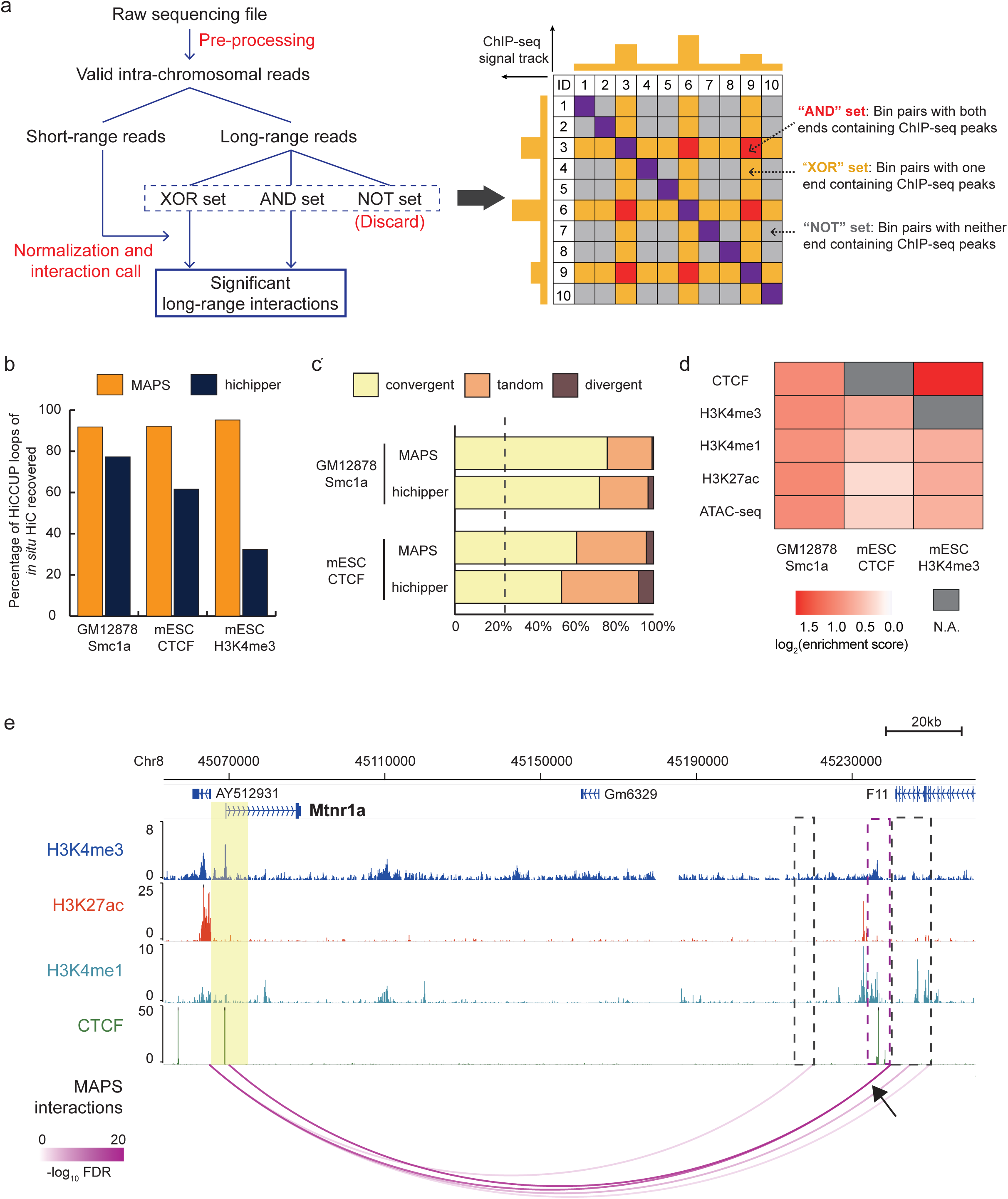
Model-based analysis of PLAC-seq and HiChIP data (MAPS). **a.** Schematic of the data analysis procedure. MAPS includes of two major components: pre-processing and interaction calling. In pre-processing, valid intra-chromosomal reads are obtained from raw sequencing file after mapping, reads pairing and duplicates removal, and then split into short-range and long-range reads depending on whether the pair of reads is separated by 1Kb or not. The long-range reads are further assigned into the “AND”, “XOR” or “NOT” set. MAPS only considers the “AND” and “XOR” set of bin pairs for normalization and identification of the statistically significant long-range chromatin interactions. Short-range reads are used to estimate and correct for biases introduced by the ChIP procedure. **b.** Comparison of sensitivity of MAPS and hichipper. The Y-axis is the sensitivity, defined as the percentage of detectable HiCCUPS loops of deeply sequenced *in situ* Hi-C data (***Table S3***) recovered by MAPS-or hichipper-identified interactions. **c.** Assessment of reliability of MAPS and hichipper. The proportion of convergent, tandem and divergent CTCF motif pairs among testable MAPS- and hichipper-identified interactions. Only interactions with both ends containing either single CTCF motif or multiple CTCF motifs in the same direction are considered. The dotted vertical line indicates the expected convergent proportion from randomly chosen CTCF motif pairs (25%). **d.** *Cis*-regulatory elements are enriched in the target bins of MAPS-identified “XOR” interactions. As only interactions from “XOR” are considered, CTCF enrichment analysis is not applicable for mESC CTCF PLAC-seq data, and H3K4me3 enrichment analysis is not applicable for mESC H3K4me3 PLAC-seq data (denoted as N.A. in the heatmap). For each ChIP-seq/ATAC-seq data, we calculated the proportion of target bins and control bins containing ChIP- seq/ATAC-seq peaks, defined as %target and %control, respectively. We further defined the enrichment score as the ratio between %target and %control. **e.** Genome-browser shows MAPS-identified interactions anchored at *Mtnr1a* promoter from mESC H3K4me3 PLAC-seq data. Anchor regions around *Mtnr1a* promoter are highlighted by yellow box (chr8:45,065,000-45,075,000, two 5Kb bins). The MAPS-identified interactions overlapping this anchor region are marked by magenta arcs. The black arrow points to the interaction verified in the previous publication^20^ and the other end of the interaction is marked by magenta box. Additional interacting regions identified by MAPS are marked by grey boxes. No interaction is identified by hichipper anchored at this region from mESC H3K4me3 PLAC-seq data.

To benchmark the performance of MAPS against hichipper, we applied both methods to three distinct PLAC-seq and HiChIP datasets, one published Smc1a HiChIP dataset from GM12878 cells^6^ and two in-house PLAC-seq datasets focusing on H3K4me3 and CTCF binding sites in the mouse embryonic stem cells (mESCs) (***Table S1***). Considering variable sequencing depth of each dataset, we used MAPS to call interactions at 5Kb resolution for GM12878 Smc1a HiChIP and mESC H3K4me3 PLAC-seq data, and at 10Kb resolution for mESC CTCF PLAC-seq data. We did not test HiCCUPS nor Fit-Hi-C, since Lareau and Aryee study^10^ has demonstrated that hichipper outperforms both HiCCUPS and Fit-Hi-C. We first examined the reproducibility of MAPS and hichipper between two biological replicates. The reproducibility of interactions called from MAPS among biological replicates ranges 69.4% ~ 85.3% for these three datasets, slightly higher than the results from hichipper (61.0% ~ 81.9% for these 3 datasets) (***Figure S4***). As a reference, using the same definition of reproducibility, the widely used HiCCUPS for *in situ* Hi-C data has 64.3% ~ 67.4% reproducibility in interaction calls between biological replicates (***Supplementary Note 2***). Since MAPS- and hichipper-identified interactions are reproducible between replicates, we combined the two biological replicates and utilized interactions identified from the combined data for all the downstream analysis. MAPS identified 37,951, 53,788 and 134,179 interacting bin pairs from GM12878 Smc1a, mESC CTCF, and mESC H3K4me3 data, respectively. The numbers of interacting bin pairs identified by MAPS are ~2-4 folds more than the numbers of hichipper (***Figure S5***). Although the median distance of the interacting bin pairs is similar between MAPS and hichipper (***Figure S5***), MAPS detects more longer-range interactions, with only 3% ~ 7% of MAPS-identified interactions spanning a distance less than 50Kb, while 16% ~ 31% of hichipper-identified interactions are within 50Kb. Noticeably, MAPS is able to identify the same set of interactions from mESC CTCF and mESC H3K4me3 data on the “common” anchors (regions with both H3K4me3 and CTCF binding), suggesting the robustness of MAPS (***Supplementary Note 3***).

Next, we investigated the sensitivity of MAPS and hichipper in detecting chromatin interactions, by comparing the results to chromatin loops identified from deeply sequenced *in situ* Hi-C data from the matching cell types^4,13^ (***Table S2***). Since PLAC-seq and HiChIP are designed to detect interactions associated with a specific protein, we filtered chromatin loops from *in situ* Hi-C data and only kept the ones associated with the protein of interest for this analysis (***Supplementary Note 4, Table S3***). MAPS achieved consistently higher sensitivity than hichipper (91.8%, 92.2%, 95.2% vs 77.3%, 61.6%, 32.5%, ***Figure 1b***) in all three datasets. The substantially improved sensitivity achieved by MAPS is partly due to the fact that MAPS can identify interactions in the “XOR” set, yielding ~2-4 folds more interactions than hichipper (***Figure S5***).

We also tried to assess the true positive rate for MAPS-and hichipper-identified interactions. However, due to lack of a gold standard of chromatin interactions in these cells, we resort to analysis of orientation of CTCF binding motifs in the identified chromatin interactions, since convergent CTCF motif has been shown to be a key feature of CTCF and cohesin-mediated chromatin loops due to loop extrusion^4, 14, 15^. Supporting the validity of the MAPS-identified interactions, convergent CTCF motif is found in 76.7%, and 61.3% testable MAPS-identified interactions from GM12878 Smc1a and mESC CTCF dataset, respectively (see ***Methods*** for details). By comparison, convergent CTCF motif is found in lower proportions of testable hichipper-identified interactions (72.8% and 53.7%, Chi-square test p-value is 3.54e-7 and <2.2e-16), suggesting that MAPS yielded more accurate calls than hichipper (***Figure 1c***). The MAPS- and hichipper-identified interactions can be grouped into singletons or clusters, depending on whether additional significant chromatin interactions exist within their neighborhoods (see ***Methods*** for details). For each interaction cluster, its summit can be further identified as the bin with the lowest FDR value. We repeated sensitivity and CTCF motif orientation analysis using only the sum of singletons and cluster summits and obtained consistent results (***Figure S6***), showing that MAPS performs almost equally well if restricted to a conservative subset of interaction calls.

One of the major advantages of MAPS over hichipper is that MAPS can detect interactions in both “AND” and “XOR” set whereas hichipper only considers the “AND” set. The “XOR” set clearly contains true positives. For example, in mESC H3K4me3 PLAC-seq data, we observe both promoter-promoter interactions (i.e., in the “AND” set) and enhancer-promoter interactions (i.e., in the “XOR” set). MAPS analysis detects many interactions in the “XOR” set in every dataset we studied (59.2% - 78.6% of MAPS-identified interactions are in the “XOR” set, see ***Table S4***), suggesting that a significant portion of MAPS-identified interactions have one anchor bin not bound by the protein of interest (hereafter referred to as the “target” bin). We then ask whether those target bins are enriched for *cis*-regulatory elements (CREs) that may contribute to gene regulation. In GM12878 Smc1a data, we found that there are twice more target bins containing ATAC-seq peaks compared to control bins (***Figure S7, Figure 1d***). This enrichment holds for H3K27ac, H3K4me1, H3K4me3 and CTCF ChIP-seq peaks (all Chi-square test p-values < 2.2e-16). We also observed enrichment of CREs among target bins in the mESC CTCF and H3K4me3 PLAC-seq datasets, suggesting the existence of functional interactions in the “XOR” set (***Figure 1d***). To further evaluate the performance between MAPS and hichipper, we examined ten experimentally determined long-range chromatin interactions in mESC^16-20^ (***Table S5***). Among them, eight were identified by MAPS using mESC H3K4me3 PLAC-seq data. By contrast, only five of them were found by hichipper (***Table S5***). In these loci, MAPS additionally identified promoter-centered long-range interactions, most of which are enriched in H3K4me1, CTCF or H3K27ac marks (***Figure 1e, Figure S8***).

A large proportion (52.6% - 87.3%) of MAPS-identified interactions do not overlap with chromatin loops identified from *in situ* Hi-C data by HiCCUPS (***Table S6***). Several lines of evidence support the reliability of these additional chromatin interactions called by MAPS. First, the “XOR” set of MAPS-specific interactions are enriched for active chromatin marks to the same degree as the chromatin interactions called by both HiCCUPS (from *in situ* Hi-C data) and MAPS (from PLAC-seq data or HiChIP datasets) (***Figure 2a, Figure S9***). Second, the MAPS-identified interactions are also detectable using SPRITE (split-pool recognition of interactions by tag extension)^21^, an orthogonal method for mapping 3D chromatin structure independent of proximity ligation. Both HiCCUPS/MAPS-shared and MAPS-specific interactions show higher interaction frequency detected by SPRITE than the matched control set, indicating that the two anchors of MAPS-specific interactions are indeed in close spatial proximity (***Figure 2b***). Third, the MAPS-identified enhancer-promoter interactions match well with functionally defined enhancer-promoter target relationships. A recent study revealed multiple functional enhancer-promoter pairs in mESCs via functional perturbation of enhancers^22^. For the five promoter-enhancer pairs spanning a distance greater than 50Kb, MAPS is able to identify three of them (***Figure 2c, Figure S10***). By contrast, none of these five enhancer-promoter pairs are defined by HiCCUPS from *in situ* Hi-C data (***Table S7***). This analysis indicates that MAPS can identify biologically relevant long-range chromatin interactions more sensitively and accurately than existing methods.

**Figure 2.**
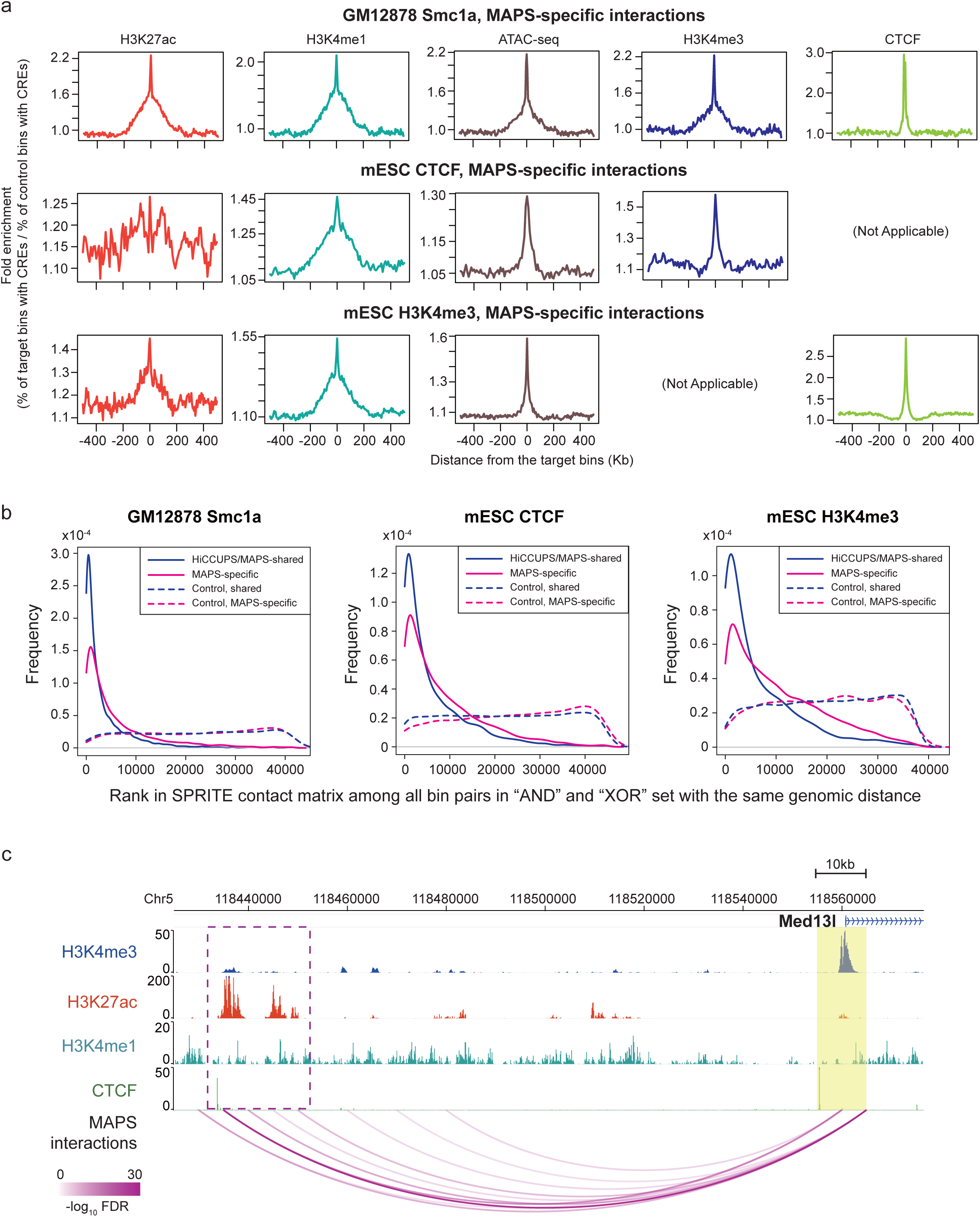
MAPS identified biologically relevant chromatin interactions from PLAC-seq and HiChIP datasets. **a.** Enrichment of CREs around the target bins of XOR set of MAPS-specific interactions. Enrichment of H3K27ac (ChIP-seq peaks), H3K4me1 (ChIP-seq peaks), ATAC-seq peaks, H3K4me3 (ChIP-seq peaks) and CTCF (ChIP-seq peaks) in a window of 500Kb around the target bins for all 3 datasets. Due to the definition of XOR set of interactions, H3K4me3 and CTCF enrichment level is not analyzed for mESC H3K4me3 and mESC CTCF PLAC-seq data, respectively. **b.** Frequency of MAPS-identified interactions and control bin pairs versus their rankings in the SPRITE contact matrix. A bin pair with higher normalized SPRITE interaction frequency tends to rank top, among all bin pairs with the same genomic distance (only bin pairs in “AND” and “XOR” sets from the SPRITE contact matrix are considered). **c.** MAPS-identified interactions from mESC H3K4me3 PLAC-seq data anchored at *Med13l* promoter. Anchor region around target promoter is highlighted by yellow box. The MAPS-identified interactions overlapping this anchor region are marked by magenta arcs. The deleted enhancer region in Moorthy et al study^22^ is marked by magenta box.

The rapidly growing popularity of PLAC-seq and HiChIP technologies necessitates the development of effective data analysis method tailored to the new datasets. MAPS takes into account of method-specific biases introduced by the ChIP procedure, and identifies long-range chromatin interactions anchored at different proteins with high reproducibility and accuracy. More importantly, MAPS can detect a large number of biologically relevant chromatin interactions that are missed by the state of art mapping approaches, making it a useful tool for investigators working on chromatin architecture, epigenomics, and gene regulatory networks.

## Data and software availability

The mESC CTCF and mESC H3K4me3 PLAC-seq data have been deposited to GEO. MAPS software can be freely downloaded from the GitHub website: https://github.com/ijuric/MAPS. (***Supplementary Note 5***).

## Acknowledgements

We would like to thank the Ren Lab members and Dr. Yin Shen for comments during the manuscript preparation. This work was supported by NIH U54DK107977 to BR and MH and R01HL129132 to YL.

## Supplementary figure legends

**Figure S1. Flowchart of MAPS pre-processing steps.** Details can be found in ***Methods*** section.

**Figure S2. Data normalization by MAPS. (a) MAPS removes biases in GM12878 Smc1a HiChIP data.** For all autosomal chromosomes, we calculated the Pearson correlation coefficients (Y-axis) between the systemic biases (effective length, GC content, mappability, ChIP efficiency) and the raw contact frequency in the “AND” set, the normalized contact frequency in the “AND” set, the raw contact frequency in the “XOR” set and the normalized contact frequency in the “XOR” set, highlighted in red, yellow, blue and purple boxes, respectively. The grey dash line presents the Pearson correlation coefficient zero. Three panels show the results in replicate 1, replicate 2, and the combined data (replicate 1 + replicate 2), respectively. **(b)** Similar to **Figure S2a**, **MAPS removes biases in mESC CTCF PLAC-seq data. (c)** Similar to **Figure S2a**, **MAPS removes biases in mESC H3K4me3 PLAC-seq data.**

**Figure S3. Contact frequency plot for chromosome 1 in mESC H3K4me3 PLAC-seq data replicate 1 at 5Kb bin resolution.** The X-axis is Log_10_ genomic distance between two interacting bins (unit: Kb). The Y-axis is the Log_10_ average raw PLAC-seq contact frequency. The red line and blue line represent the contact probability for bin pairs in the “AND” and “XOR” set, respectively.

**Figure S4. The reproducibility of MAPS- and hichipper-identified interactions between two biological replicates.** The Y-axis represents the percentage of reproducible interactions among two biological replicates. “rep1” denotes the proportion of interactions identified in biological replicate 1 reproducible in biological replicate 2, whereas “rep2” denotes the proportion of interactions identified in biological replicate 2 reproducible in biological replicate 1.

**Figure S5. Summary of MAPS-and hichipper-identified interactions of all three datasets. (a) The number of interactions and the distribution of interaction length from MAPS-identified interactions.** The left, middle and right column represents the MAPS calls from GM12878 Smc1a HiChIP, mESC CTCF PLAC-seq and mESC H3K4me3 PLAC-seq combined data (replicate 1 + replicate 2), respectively. Each histogram shows the distribution of interaction length. The vertical blue bar represents the median distance of interactions. **(b)** Similar to **Figure S5a**, **the number of interactions and the distribution of interaction length from hichipper-identified interactions.**

**Figure S6. Sensitivity and convergent CTCF motif analysis results using only singletons and cluster summits from MAPS- and hichipper-identified interactions. (a) Similar to Figure 1b, the sensitivity of MAPS- and hichipper-identified interactions, using only singletons and cluster summits. (b) Similar to Figure 1c, the proportion of convergent CTCF motif pairs among testable MAPS- and hichipper-identified interactions, using only singletons and cluster summits.** The dotted vertical line indicates the expected convergent proportion from randomly chosen CTCF motif pairs (25%).

**Figure S7. Cartoon illustration of anchor bin, target bin and control bin used in cis-regulatory elements enrichment analysis (related to Figure 1e, Figure 2a and Figure S9).** The solid yellow curve at left represents a statistically significant long-range chromatin interaction in the “XOR” set, connecting the anchor bin (the red box) and the target bin (the blue box). The dashed yellow curve at right represents a random collision between the anchor bin (the red box) and the control bin (the purple box). The interaction and the random collision has the same genomic distance.

**Figure S8. MAPS-identified interactions from mESC H3K4me3 PLAC-seq data anchored at: (a) *Pou5f1* promoter, (b) *Sox2* promoter, (c) *Tbx5* promoter, (d) *Wnt6* promoter, (e) *Nanog* promoter.** Anchor regions around target promoter are highlighted by yellow boxes. The MAPS-identified interactions overlapping the anchor regions are marked by magenta arcs. The black arrow points to the interaction verified in previous publications16-20 and the other end of the interaction is marked by magenta boxes. Additional interacting regions identified by MAPS are marked by grey boxes.

**Figure S9. Enrichment of CREs around the target bins of XOR set of HiCCUPS/MAPS-shared interactions.** Enrichment of H3K27ac (ChIP-seq peaks), H3K4me1 (ChIP-seq peaks), ATAC-seq peaks, H3K4me3 (ChIP-seq peaks) and CTCF (ChIP-seq peaks) in a window of 500Kb around the target bins for all 3 datasets. Due to the definition of XOR set of interactions, H3K4me3 and CTCF enrichment level is not analyzed for mESC H3K4me3 and mESC CTCF PLAC-seq data, respectively.

**Figure S10. MAPS-identified interactions from mESC H3K4me3 PLAC-seq data anchored at: (a) *Elt4* promoter (chr2:20,515,000-20,525,000), (b) *Ifitm3* promoter (chr7:141,005,000-141,015,000).** Anchor regions around target promoters are highlighted by yellow boxes. The MAPS-identified interactions overlapping this anchor region are marked by magenta arcs. The deleted enhancer regions in Moorthy et al study^22^ are marked by magenta boxes.

**Figure S11. MAPS running time (Y-axis) increases linearly proportional to overall sequencing depth (X-axis).**

## Supplementary table titles

**Table S1. A brief summary of three PLAC-seq and HiChIP datasets used in this study.**

**Table S2. A brief summary of HiCCUPS loops identified from deeply sequenced in situ Hi-C data.**

**Table S3. Number of HiCCUPS loops which are detectable in each PLAC-seq and HiChIP data.**

**Table S4. Number of MAPS-identified interactions in AND and XOR sets.**

**Table S5. Ten long-range promoter-centered interactions in mESC verified by 3C or 4C in previous publications**^16-20^.

**Table S6. Overlap between MAPS-identified interactions and HiCCUPS loops in all three PLAC-seq and HiChIP datasets.**

**Table S7. A list of functional validated enhancer-promoter pairs in mESC from Moorthy et al study^22^. Only enhancer-promoter pairs are >50Kb in genomic distance are listed.**

**Table S8. Summary of all sequencing data sets used in this study.**

## Methods

### Sequencing Data

All data sets used (both external and in-house generated for this study) are summarized in *Table S8*.

### Cell culture and fixation

The *F1 Mus musculus castaneus* × S129/SvJae mouse ESC line (F123 line) was a gift from Dr. Rudolf Jaenisch and was previously described^23^. F123 cells were cultured in DMEM (10013-CV, Corning), supplement with 15% knockout serum replacement (10828028, Invitrogen), 1×penicillin/streptomycin (15140122, Thermo Fisher Scientific), 1×non-essential amino acids (11140050, Thermo Fisher Scientific), 1×GlutaMax (35050061, Thermo Fisher Scientific), 1000 U/ml LIF (ESG1107, Millipore), 0.1 mM β-mercaptoethanol (M3128, Sigma). F123 cells were maintained on irradiated CF1 mouse embryonic fibroblasts (A34180, Thermo Fisher Scientific) and were passaged once on 0.1% gelatin-coated feeder-free plates before harvesting.

Cells were harvested by accutase treatment and resuspended in culture medium described above but without knockout serum replacement at a concentration of 1×10^6^ cells per 1ml. Methanol-free formaldehyde solution was added to a final concentration of 1% (v/v) and fixation was performed at room temperature for 15 min with slow rotation. The fixation was quenched by addition of 2.5 M glycine solution to a final concentration of 0.2 M with slow rotation at room temperature for 5 min. Fixed cells were pelleted by centrifugation at 2,500×*g* for 5 min at 4 °C and washed with ice-cold PBS once. The washed cells were pelleted again by centrifugation, snap-frozen in liquid nitrogen and stored at −80 °C.

### PLAC-seq on F123 cells

PLAC-seq libraries were prepared using method as previously described^5^. The detailed experimental procedures are provided in ***Supplementary Note 7***. In brief, 1-3 million crosslinked F123 cells were digested 2 hours at 37 °C using 100 U MboI followed by biotin fill-in and proximity ligation at room temperature for 4 hours. Then the nuclei were further lysed, sonicated and immunoprecipitated against the antibodies of choice. After immunoprecipitation, reverse crosslink was performed overnight at 65 °C after adding proteinase K to extract DNA. DNA fragments containing ligation junctions were enriched with streptavidin beads followed by on-beads end repair, A-tail adding, adapter ligation and PCR amplification for 12-13 cycles.

### ATAC-seq on F123 cells

ATAC-seq was performed using method as previously described^24^. In brief, 100,000 freshly harvested F123 cells were resuspend in lysis buffer (10 mM Tris-HCl, pH 7.4, 10 mM NaCl, 3 mM MgCl_2_ and 0.1% IGEPAL CA-630) and rotate at 4°C for 15 minutes. After lysis the nuclei was spun down at 500×*g* for 5 min at 4 °C. Then the reaction was carried out for 30 min at 37 °C in 1×TD buffer with 2.5 μL transposase from Nextera DNA Library Prep Kit (Illumina). After reaction completion DNA is purified using MinElute PCR Purification Kit (Qiagen). PCR amplification was perform with 1×NEBNext PCR MasterMix and 1 μM i7-index and i5-index primers using the following PCR condition: 72 °C for 5 min; 98 °C for 30 s; and 8 cycles of 98 °C for 10 s, 63 °C for 30 s and 72 °C for 1 min. The amplified libraries are purified and size selected using 0.55× and 1.5× (total) of sample volume.

### ChIP-seq on F123 cells

2 million fixed F123 cells were thawed on ice, resuspend in hypotonic lysis buffer (20 mM HEPES, pH 8.0, 10 mM KCl, 1 mM EDTA, 10% glycerol) with proteinase inhibitors and rotate at 4 °C for 15 minutes. The nuclei were then washed once with hypotonic lysis buffer with proteinase inhibitors and resuspend in 130 μL RIPA buffer (10 mM Tris, pH 8.0, 140 mM NaCl, 1 mM EDTA, 1% Triton X-100, 0.1% SDS, 0.1% sodium deoxycholate) with proteinase inhibitors. After incubation on ice for 10 minutes, the nuclei were sheared using Covaris M220 with following setting: power, 75 W; duty factor, 10%; cycle per burst, 200; time, 10 minutes; temp, 7 °C. The cell lysate was cleared by centrifugation at 15,000×*g* for 20 min and supernatant was collected. The clear cell lysate was precleared with Protein G Sepharose beads (GE Healthcare) and for 3 hours at 4 °C with slow rotation. ~5% of precleared cell lysate was saved as input control. The rest of the lysate was mixed with 2.5 μg of H3K4me3 (04-745, Millipore) antibody and rotate at 4 °C for at least 12 hours. On the next day, 0.5% BSA-blocked Protein G Sepharose beads (prepared one day ahead) were added and rotated for another 3 hours at 4 °C. The beads were collected by centrifugation at 400×*g* for 1 min and then washed with RIPA buffer three times, high-salt RIPA buffer (10 mM Tris, pH 8.0, 300 mM NaCl, 1 mM EDTA, 1% Triton X-100, 0.1% SDS, 0.1% sodium deoxycholate) twice, LiCl buffer (10 mM Tris, pH 8.0, 250 mM LiCl, 1 mM EDTA, 0.5% IGEPAL CA-630, 0.1% sodium deoxycholate) once, TE buffer (10 mM Tris, pH 8.0, 0.1 mM EDTA) twice. Washed beads were treated with 10 μg Rnase A in extraction buffer (10 mM Tris, pH 8.0, 350 mM NaCl, 0.1 mM EDTA, 1% SDS) for 1 hours at 37 °C, followed by reverse crosslinking in the presence of proteinase K (20 μg) overnight at 65 °C. After reverse crosslink the DNA was purified by Zymo DNA Clean&Concentrator. For library preparation, 10-100 ng ChIP DNA or input DNA was first end repaired at 20 °C for 30 minutes in 1×T4 DNA ligase buffer (NEB) with 0.5mM dNTP mix, 3U T4 DNA polymerase (NEB), 2.5U Klenow fragment (NEB) and 10U T4 PNK (NEB). The repaired DNA was then purified by Zymo DNA Clean&Concentrator and adenylated at 37 °C for 30 minutes in 1×NEBbuffer 2 (NEB) with 0.4mM dATP, 10U Klenow fragment (3’-5’ exo-) (NEB). The adenylated was purified by Zymo DNA Clean&Concentrator and ligated to the adapters (Illumina, TruSeq, 0.1 μL per 100ng DNA) at 16 °C for overnight in 1×T4 DNA ligase buffer (NEB) with 400U T4 DNA ligase. After purification with Zymo DNA Clean&Concentrator, DNA was amplified with KAPA HiFi HotStart ReadyMix PCR Kit for 12 cycles according to the manufacturer’s instructions. The amplified libraries were purified with Ampure Beads to extract fragments between 200-600bp for sequencing.

### ChIP-seq data processing

The H3K4me3 ChIP-seq data on F123 cells was analyzed using ENCODE Uniform processing pipeline for ChIP-seq (histone marks) (https://github.com/ENCODE-DCC/chip-seq-pipeline) with default parameters.

### ATAC-seq data processing

ATAC-seq reads were mapped to mm10 genome using bowtie 1.1.2 with flags “-X2000 --no-mixed --no-discordant”. The reads were converted to bam files, sorted, and PCR duplicates and mitochondrial reads were removed using samtools. To account for the Tn5 insertion position, read end positions were moved 4bp towards the center of the fragment. Bigwig signal tracks and peak calls were generated using MACS2 2.1.1.20160309 and the following flags: “-nomodel-shift 37-ext 73-pval 1e-2 -B- SPMR-call-summits”. To obtain the set of replicated peaks for each sample, the data were processed as described above for each replicate independently as well as pooled. Using bedtools 2.27.1, the pooled peaks were intersected against each replicate’s peaks sequentially, and pooled peaks present in both replicates were considered to be ‘replicated’.

### MAPS pre-processing component

MAPS takes the raw paired-end reads (fastq files) from PLAC-seq and HiChIP experiment as input, maps them to the reference genome (***Figure S1***). Specifically, we use “bwa mem” to map each end of paired-end reads to the reference genome separately (mm10 or hg19, ***Table S1***), and remove non-mappable reads and low mapping quality reads. We further remove read pairs mapped to less than 2 or more than 3 locations. The read pairs mapped to only 1 location contain no information about chromatin structures whereas the chance of a pair mapped to more than 3 locations are rare and such pairs most likely represent spurious ligation events. A read pair is defined as “valid” if it can be mapped to exactly 2 locations. For a read pair mapped to 3 locations, if all 3 locations are on the same chromosome, it spans over one ligation junction. In this case, the mapping pair with the second largest linear distance is defined as “valid”, since it represents the pair that is closer to the ligation junction. If 3 locations are on 2 different chromosomes, in most cases the 2 locations within the same chromosome are close to each other, therefore we randomly select one of the two locations on the same chromosome, and pair with the location on the other chromosome. The chance of 3 locations are on 3 different chromosomes is very low and such pairs most likely represent spurious ligation events, therefore we discard such pairs. After pairing all the reads as described above, we use “samtools rmdup” to remove PCR duplicates. Furthermore, we select short-range reads (<=1Kb) with two reads from different strands to measure IP efficiency, and keep long-range reads (>1Kb) to identify long-range chromatin interactions. MAPS then extracts the intra-chromosomal long-range reads and takes the ChIP-seq peaks of protein of interest as the interaction anchors (***Table S8***), and groups all bin pairs into the “AND”, “XOR” and “NOT” sets. MAPS only selects the “AND” and “XOR” sets for the next data normalization step. When the ChIP-seq data is not available, one can apply hichipper to PLAC-seq and HiChIP data to obtain interaction anchors.

Data normalization is a challenging issue for any chromatin interaction data. Notably, the matrix-balancing algorithms used for Hi-C data normalization, including ICE^25^, VC^26^ and KR^4^, are inappropriate for PLAC-seq and HiChIP data normalization. Due to the ChIP procedure, in theory the bins with protein binding always have much higher total number of contacts compared to the bins without protein binding, which violates the crucial “equal visibility” assumption implied by the matrix-balancing algorithms.

To accommodate unique features of PLAC-seq and HiChIP data, we propose to extend our previous HiCNorm^12^ method to normalize PLAC-seq and HiChIP data. Let *x*_*ji*_ represent the read count (i.e., number of paired-end reads) spanning between bin *i* and bin *j*. Due to symmetry, we only consider bin pairs (*i,j*) with *i*< *j*. In addition, we only consider intra-chromosomal contacts within 1Mb, and do not use two adjacent bin pairs (*j*− *i*≥ 2). Let *f*_*i*_,*gc*_*i*_,*m*_*i*_ and *IP*_i_ represent the effective fragment length, GC content, mappability score, and antibody IP efficiency (measured by the number of short-range reads, i.e., intra-chromosomal reads <=1Kb) of bin *i*, respectively. The definition of *f*_*i*_,*gc*_*i*_ and *m*_*i*_ are described in HiCNorm^12^. Also, let *d*_*ij*_ denote the genomic distance between bin *i* and bin *j*. We only model bin pairs (*i,j*) with non-zero count (*x*_*ji*_ ≥ 1), and assume that *x*_*ji*_ follows a zero-truncated Poisson (ZTP) distribution with mean *μ*_*ji*_, where

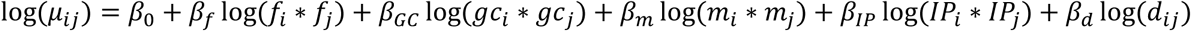

Here *β*_*0*_ is the intercept for overall sequencing depth. *β*_*f*_, *β*_*GC*_, *β*_*m*_, *β* _*IP*_ and *β* _*d*_ are regression coefficients for effective fragment length, GC content, mappability score, antibody IP efficiency and genomic distance, respectively. We fit the aforementioned ZTP regression model for each chromosome, separately for the “AND” set and the “XOR” set, using R function “ppois” in the “VGAM” library, to obtain the expected contacted frequency *e*_*ij*_ for each bin pair (*i,j*). These *e*_*ij*_ ‘s represent background expected from random chromatin collisions. Next, we calculate a ZTP p-value for each bin pair (*i,j*), defined as *p*_*ij*_ = ZTP(X> *x*_*ij*_ | *e*_*ij*_). Similar to Fit-Hi-C, we view bin pairs with extremely low p-values (< 1 / total number of non-zero bin pairs) as outlies. We then remove those outliers, and re-fit the ZTP regression model using the remaining data to re-calibrate the background, obtaining re-calibrated expected contact frequency 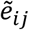 and corresponding ZTP p-value 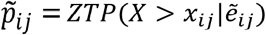 We further convert ZTP p-value 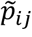 into false discovery rate (FDR) *q*_*ij*_ using R function “p.adjust”. Within each chromosome, the FDR is calculated by the “AND” and “XOR” set, separately.

### MAPS interaction calling component

We then identify statistically significant long-range chromatin interactions from normalized PLAC-seq and HiChIP data. Specifically, we define a bin pair (*i,j*) as an statistically significant bin pair if it satisfies the following three criteria simultaneously: (1) *x*_*ij*_ ≥ 12, 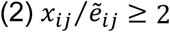 and (3) *q*_*ij*_ < 0.01. Details of justification of such thresholds can be found in ***Supplementary Note 6***. Starting from these significant bin pairs, we further group adjacent ones into clusters, and singletons (defined as isolated significant bin pairs without adjacent ones). Specifically, we denote *d*_*ij*_ as the genomic distance between bin *i* and bin *j*, and group significant bin pair (*i,j*) and significant bin pair (*m,n*) into the same interaction cluster if max{*d*_*im*_, *d*_*nj*_} ≤ 15Kb. Each significant bin pair belongs to one unique cluster, or it is a singleton. For the significant bin pairs defined as singletons, we apply additional filtering and only keep the ones with *q*_*ij*_ < 10^−4^ as significant interactions since singletons are more likely to be false positives. For the significant bin pairs as part of a cluster, we keep all of them as significant interactions. For each interaction cluster, we further identify its summit, defined as the bin pair(s) with the lowest FDR. Therefore, the final MAPS output contains the following information: 1) a list of statistically significant long-range chromatin interactions; 2) for each interaction, whether it is a singleton or belongs to a cluster; 3) if an interaction is part of a cluster, whether it is the summit of this cluster and which interactions are in the same cluster.

### Identification of interactions with hichipper

To call interactions from the same PLAC-seq and HiChIP datasets using hichipper, we performed the mapping and preprocessing using the default settings of HiC-Pro 2.7.6 and bowtie 2.3.0 as base mapper (recommended by hichipper), specifying digestion fragment size of 100 to 100,000. Genome fragment size files were obtained from the GitHub repository of hichipper (https://github.com/aryeelab/hichipper). Since the data in each run are from one sample and no merging was required, therefore we removed allValidPairs and mRStat files to make the HiC-Pro output consistent with the requirements of the hichipper input. We then used hichipper v0.4.4 to call interactions using ChIP-seq peaks as interaction anchors (***Table S8***). In the default setting, hichipper outputs all interacting bin pair with *x*_*ij*_ ≥ 2. Such liberal threshold leads to a large number of false positives (data not shown). To make a fair comparison with MAPS, we used the same thresholds described above for MAPS calls to filter the outputs from hichipper, only keeping hichipper-identified interactions with *x*_*ij*_ ≥ 12 and *q*_*ij*_ < 0.01 (hichipper does not output the expected contact frequency, so the filter based on normalized contact frequency cannot be applied). We then group hichipper-identified interactions into interaction clusters and singletons using the same definition as described above and further filter the interactions defined as singletons using FDR threshold *q*_*ij*_ < 10^−4^. The final hichipper output has the same format as the MAPS output.

### HiCCUPS loops from *in situ* Hi-C data

The HiCCUPS loops of GM12878 are acquired from Rao et al. study4. Specifically, file “GSE63525_GM12878_primary+replicate_HiCCUPS_looplist.txt.gz” was downloaded, which contains in total 9,448 loops. Among these 9,448 loops, we selected 6,316 loops where both two interacting anchors are 5Kb bins (***Table S2***). To generate the 5Kb and 10Kb resolution of HiCCUPS loops of mESCs, we downloaded the raw fastq files of all four biological replicates from Bonev et al. study^13^ and performed mapping, pairing reads and PCR duplicates removal in the same way as we did for PLAC-seq and HiChIP data (refer to “**MAPS pre-processing component”** above). Afterwards we combined the valid pairs from all four replicates and then applied HiCCUPS to call loops at 5Kb and 10Kb resolution with the following parameters: “-r 5000,10000-k KR-f .1,.1 -p 4,2 -i 7,5 -t 0.02,1.5,1.75,2 -d 20000,20000” (***Table S2***).

### Reproducibility analysis

We evaluate the reproducibility of MAPS- and hichipper-identified interactions between two biological replicates. We denote *d*_*ij*_ as the genomic distance between bin *i* and bin *j*. We then define an interaction, bin pair (*i,j*), in one replicate is reproducible, if and only if there exists an interaction, bin pair (*m,n*), in the other replicate such that max{*d*_*im*_,*d*_*ij*_} ≤ 15Kb.

### Sensitivity analysis

We evaluate the sensitivity of MAPS- and hichipper-identified interactions, using HiCCUPS loops called from deeply sequenced *in situ* Hi-C datasets as true positives. Specifically, we used GM12878 *in situ* Hi-C data with ~4.9 billion reads from Rao et al. study^4^, and mESC *in situ* Hi-C data with ~7.3 billion reads from Bonev et al. study^13^. We first selected a subset of HiCCUPS loops which are detectable in corresponding PLAC-seq and HiChIP data (***Table S3***, ***Supplementary Note 4***). Next, we defined a HiCCUPS loops, bin pair (*i,j*), is re-discovered from PLAC-seq and HiChIP data, if and only if there exists an interaction, bin pair (*m,n*), in PLAC-seq and HiChIP data such that max{*d*_*im*_,*d*_*ij*_} ≤ 15Kb. The sensitivity is calculated by the ratio between the number of HiCCUPS loops re-discovered from PLAC-seq and HiChIP data and the total number of HiCCUPS loops detectable in PLAC-seq and HiChIP data.

### CTCF motif orientation analysis

We examine the CTCF motif orientation of testable MAPS- and hichipper-identified interactions. Specifically, we first download the CTCF ChIP-seq peak lists of GM12878 and mESC (***Table S8***) and then search for all the CTCF sequence motifs among those peak using FIMO^27^ (default parameters) and the CTCF motif (MA0139.1) from the JASPAR^28^ database. Based on this list of CTCF motifs, we then select a subset of MAPS- or hichipper-identified interactions with both ends containing either single CTCF motif or multiple CTCF motifs in the same direction. Finally, we count the frequency of four possible directionality of CTCF motif pairs, and calculate the proportion of convergent, tandem and divergent CTCF motif pairs among all testable interactions.

### *Cis*-regulatory elements enrichment analysis

For two interacting bins in the “XOR” set, we define the bin which binds to the protein of interest as the “anchor” bin, and the bin which does not bind to the protein of interest as the “target” bin. In order to access the biological relevance of peaks in the “XOR” set, we evaluated whether *cis*-regulatory elements are enriched within those target bins, compared to the control bins which are in the same distance with the anchor bin, but do not bind to the protein of interest (***Figure S7***).

The ChIP-seq and ATAC-seq data used for this analysis is summarized in ***Table S8***. For each ChIP-seq/ATAC-seq data, we calculated the proportion of target bins and controls containing ChIP-seq/ATAC-seq peaks, defined as %target and %control, respectively. We further defined the enrichment score as the ratio between %target and %control.

### Definition of MAPS-specific interactions and HiCCUPS/MAPS-shared interactions

We divided all MAPS-identified interactions into two groups based on their overlap with HiCCUPS loops. Similar to our method in the sensitivity analysis, we defined a MAPS interaction, bin pair (*i,j*), is overlapped with a HiCCUPS loop, if and only if there exists an interaction, bin pair (*m,n*), in HiCCUPS loop list such that max{*d*_*im*_, *d*_*ij*_} ≤ 15Kb. If a bin pair (*i,j*) is overlapped with a HiCCUPS loop, we define it as a HiCCUPS/MAPS-shared interaction. If a bin pair (*i,j*) is not overlapped with a HiCCUPS loop, we define it as a MAPS-specific interaction. The number of MAPS-specific interactions and HiCCUPS/MAPS-shared interactions are listed in ***Table S6***.

### Validation of MAPS-specific interactions by SPRITE data

The normalized SPRITE interaction frequency matrices were downloaded from GEO with access number GSE114242^21^. The GM12878 and mESC SPRITE data is at 25Kb bin and 20Kb bin resolution, with reference genome hg19 and mm9, respectively. The MAPS-identified interactions consist of two types of bin pairs: singletons and interaction clusters (defined in “**MAPS interaction calling component**” section). For each singleton bin pair (*i,j*), we define it as “shared” if there exists a bin pair (*m,n*) in HiCCUPS loop such that max{d_im_, d_ij_} ≤ 15Kb. Otherwise, we define the singleton bin pair (*i,j*) as “MAPS-specific”. For each interaction cluster, we define it as “shared” if any one bin pair in the interaction cluster is a “shared” bin pair. Otherwise, if all bin pairs in an interaction cluster are “MAPS-specific”, we define the entire interaction cluster as “MAPS-specific”. Since PLAC-seq and HiChIP data is at higher resolution than SPRITE data, we only selected singletons and the summits of “shared” and “MAPS-specific” interaction clusters for the downstream analysis. We then zoomed the selected bin pairs out to the matched lower resolution in SPRITE data for a fair comparison. Specifically, for MAPS-identified interactions from GM12878 Smc1a HiChIP data, we first selected the center position of 5Kb interacting bin of an interaction summit, and then allocated the 25Kb bin containing that center position. This procedure created a list of 25Kb bin pair, among which each contains MAPS-identified interaction summit. Similarly, for MAPS-identified interactions from mESC CTCF and mESC H3K4me3 PLAC-seq data, we first selected the center position of 10Kb/5Kb interacting bin of an interaction summit (reference genome mm10), converted it into reference genome mm9 using UCSC Liftover tool (https://genome.ucsc.edu/cgi-bin/hgLiftOver), and then allocated the 20Kb bin containing that center position. This procedure created a list of 20Kb bin pair, among which each contains MAPS-identified interaction summit.

To evaluate the normalized SPRITE interaction frequency for MAPS-identified interactions, we used the following procedure to create the control set. For a bin pair in the “XOR” set, we defined the bin with ChIP-seq peak as the “anchor” bin and the bin without ChIP-seq peak as the “target” bin. We then find the “control” bin and such as the “anchor” bin has the same genomic distance between the “target” bin and the “control” bin (***Figure S7***). The control bin pair is defined as the pair of the “anchor” bin and the “control” bin. For a bin pair in the “AND” set, since both two bins contain ChIP-seq peak, we randomly selected one bin as the “anchor” bin, and defined the remaining one as the “target” bin. Next, we repeated the procedures described above to find the “control” bin, and created the control bin pair for the bin pairs in the “AND” set. Finally, we filtered out any control bin pairs which are overlapped with MAPS-identified interactions. Let *S*_*ij*_ represent the normalized SPRITE interaction frequency between 25Kb/20Kb bin *i* and *j*. We defined the rank of bin pair (*i, j* as the number of bin pair in the same genomic distance, but with higher normalized SPRITE interaction frequency than*S*_*ij*_. Here in the normalized SPRITE interaction frequency matrices, we only used all bin pairs in the “AND” and “XOR” set.

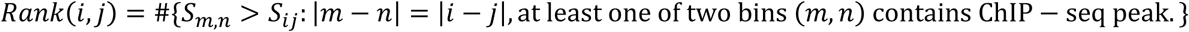

A bin pair with higher normalized SPRITE interaction frequency tends to rank top, among all bin pairs with the same genomic distance.

